# Growth and grain yield of eight maize hybrids is aligned with water transport, stomatal conductance, and photosynthesis in a semi-arid irrigated system

**DOI:** 10.1101/2021.02.04.429760

**Authors:** Sean M. Gleason, Lauren Nalezny, Cameron Hunter, Robert Bensen, Satya Chintamanani, Louise H. Comas

## Abstract

There is increasing interest in understanding how trait networks can be manipulated to improve the performance of crop species. Working towards this goal, we have identified key traits linking the acquisition of water, the transport of water to the sites of evaporation and photosynthesis, stomatal conductance, and growth across eight maize hybrid lines grown under well-watered and water-limiting conditions in Northern Colorado. Under well-watered conditions, well-performing hybrids exhibited high leaf-specific conductance, low operating water potentials, high rates of midday stomatal conductance, high rates of net CO_2_ assimilation, greater leaf osmotic adjustment, and higher end-of-season growth and grain yield. This trait network was similar under water-limited conditions with the notable exception that linkages between water transport, midday stomatal conductance, and growth were even stronger than under fully-watered conditions. The results of this experiment suggest that similar trait networks might confer improved performance under contrasting climate and soil conditions, and that efforts to improve the performance of crop species could possibly benefit by considering the water transport pathway within leaves, as well as within the whole-xylem, in addition to root-level and leaf-level traits.

## Introduction

Crop performance is an outcome of the coordinated functioning of many physiological processes. The need for a holistic understanding of the mechanisms underlying plant performance is becoming increasingly recognized, especially for complex responses such as growth and grain production under drought (Tardieu et al. 2018). Crop improvement could likely be facilitated by considering multiple physiological traits together, as well as the connections between these traits and how these connections shift under different soil and climate scenarios (Gleason et al. 2019). Although the idea that selection for multiple traits might result in better outcomes has been suggested previously (Campos et al. 2004; Condon 2020), what is becoming more clear is the need to include linkages connecting soil water, its transport to (near) the stomata, and the photochemistry that these processes support (Turner et al. 2014; Brodribb et al. 2015; Gleason et al. 2017a). Several physiological traits have been found to affect crop performance when studied in isolation of one another. For example, root (Comas et al. 2013; White 2019), xylem (Tombesi et al. 2010; Ryu et al. 2016; Gleason et al. 2017a; Cardoso et al. 2018), stomatal regulation (Zaman-Allah et al. 2011; Messina et al. 2015), and photochemistry (Rocher et al. 1989; Galic et al. 2019), have all been shown to influence water use, water use efficiency, and growth. Given the efficacy of each of these traits to affect plant functioning, as well as the clear and well-understood physiological linkages among them, it is possible that crop performance could be improved by selecting for specific trait combinations matched with different climate and soil scenarios.

Here, we focus on traits conferring improved water transport, and “drought resistance”, i.e., the ability of a plant to maintain growth and reproductive fitness when the soil and xylem water potentials are low (Passioura 2006; Volaire 2018). Linkages among aridity (atmosphere and soil), leaf area, hydraulic conductance, and photosynthesis, can be understood via the Penman-Monteith equation and Darcy’s law, as modified by Whitehead and Jarvis (Whitehead et al. 1984; Whitehead 1998) (hereafter the Whitehead and Jarvis proportionality):

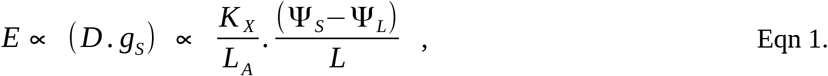

where E = transpiration, D = the leaf-to-atmosphere vapor pressure deficit, g_S_ = stomatal conductance, K_x_ = xylem-specific conductivity, L_A_ = leaf area, (Ψ_S_ - Ψ_L_) = water potential gradient between soil and leaf, and L = path-length between soil water and the sites of evaporation within the leaf.

Eqn. 1 represents an approximation of how we might expect leaf area, xylem-specific conductance, and the driving force (pressure gradient) to relate to one another (Whitehead et al. 1984). For example, if we accept that CO_2_ must pass through the stomata before it can be “fixed” by either pep-carboxylase (C4) or rubisco (C3, C4), then we must also accept that water will pass out the stomata as a consequence (i.e., the left side of the proportionality) (Whitehead 1998). This water that is spent to obtain CO_2_ must be delivered to the stomata via the vasculature (K_X_) (Brodribb et al. 2007). If we wish to double the stomatal conductance of a given plant or leaf, then we must also double K_X_, or the driving force (Ψ_S_ - Ψ_L_), or decrease either L_A_ or L by one half (Gleason et al. 2012). Each of these “choices” comes with a cost/risk, the magnitude of which depends on the climate, soil, and competitive interactions with neighboring plants.

It is clear from Eqn. 1 that xylem-specific conductivity (K_X_) is well-positioned to provide hydraulic balance in the face of greater atmospheric and/or soil aridity, albeit with added investment in vasculature (Gleason et al. 2012, 2019). This may explain why xylem-specific conductivity (K_X_) varies enormously across species and habitats (nearly three orders of magnitude), far more than any other trait in the Whitehead-Jarvis proportionality (He et al. 2019; Liu et al. 2019). Xylem-specific conductivity, as well as the susceptibility of the xylem to failure, have recently been reported as important traits conferring drought resistance in monocotyledon crop species (Guha et al. 2018; Wang et al. 2018; Gleason et al. 2019). A such, it has also been suggested that efforts to improve crop performance in drought-prone environments might benefit by explicitly considering xylem traits and water transport between the soil and leaf (Brodribb et al. 2015; Gleason 2015).

There are, of course, other traits that confer improved performance under limited water availability. For example, much research over the last two decades has focused on the improvement of transpiration efficiency, either via higher photosynthesis (preferred) or reduced transpiration (less desirable) (Zhu et al. 2010; Gilbert et al. 2011; Messina et al. 2015; Sinclair 2018). This can be understood in the context of Eqn. 1, as carbon income per unit water that has been invested (“E”; left side of the equation) to obtain this carbon, i.e., the seasonally-integrated CO_2_ ~ H_2_O exchange rate. The efficacy of these traits to confer better performance under limited water availability are supported by sound theoretical constructs, and should be most effective in environments where either much of the received precipitation can be passed through crop stomata (in exchange for CO_2_), or that precipitation received early in the season can be “banked” in the soil and used conservatively until it is needed later in the season (e.g., during anthesis). This strategy, and traits aligned with it, have been discussed at length elsewhere (Turner et al. 2014; Vadez et al. 2014; Sinclair 2018) and therefore will not be discussed further here, however, we note that it is important to realize that traits conferring soil water extraction and/or transport may in some cases be incompatible with traits conferring higher transpiration efficiency (Blum 2009; Turner et al. 2014). For example, if precipitation received early in the season is needed later in the season, conservative stomatal behavior and reduced transpiration may be preferable to higher stomatal conductance and low operating water potentials (Vadez et al. 2014; Sinclair et al. 2017). As such, the experiment described here should be considered carefully in the context of the soil and climate characteristics of the study site, and importantly, we might expect different trait combinations to confer improved performance under different soil and climate conditions (Tardieu et al. 2018). Specifically, is necessary to evaluate the efficacy of different trait combinations in the context of seasonal precipitation patterns, antecedent soil water, and other competing water “sinks”, e.g., evaporation from the soil surface, saturated and unsaturated movement in soil beyond the reach of the roots, and soil water uptake by weeds.

We examined the efficacy of using conceptual and quantitative trait networks as tools to understand the linkages between water, carbon income, and grain production across eight maize hybrids grown in the sami-arid environment of the Colorado High Plains. We applied a quantitative framework (the Whitehead and Jarvis proportionality) to help us choose which physiological traits were most likely to affect water extraction, water transport, and water use efficiency under water-limited and non-limited scenarios. We addressed the following questions: 1) Are biomass increment and grain yield dependent on water conductance traits (xylem, leaf, stomata), 2) are traits that are necessary for improving growth via water conductance and use (leaf hydraulic conductance, operating leaf water potential, maximal stomatal conductance, midday stomatal conductance, and CO_2_ assimilation) operating as a connected network, i.e., is there meaningful covariation among these traits and is this covariation logical (i.e., strength and direction), and 3) are there specific traits and trait linkages which appear to be good targets for crop improvement programs?

## Materials and methods

### Site and hybrid selection

This experiment was conducted at USDA’s Limited Irrigation Research Farm near Greeley, Colorado, USA (40.4486° latitude, −104.6368° longitude). The mean monthly minimum temperature during the growth season (May – October, 2017) was 8.4 °C, mean monthly maximum temperature was 25.9 °C, and mean monthly precipitation was 4.9 cm (Colorado Agricultural Meteorological Network 2020). Soils on the site range from sandy loam to clay loam (Ustic Haplargids). Maize (*Zea mays* L.) hybrids were chosen to represent a wide range of drought tolerance from experimental trials performed in La Salle, Colorado (Syngenta AG, Basel Switzerland).

### Experimental design

All maize hybrids were grown under fully-watered (hereafter “wet”) and water deficit (hereafter “dry”) treatments. Wet and dry treatments were designed to deliver either 100% or 40% of the evapotranspiration measured on a reference maize hybrid. Plots were watered once each week via drip irrigation to maintain these target ET levels. Thus, all plots (hybrids) within each irrigation treatment (40% or 100% ET) received the same amount of water during each irrigation, i.e. irrigation water was not adjusted to account for differences in transpiration/evaporation among hybrids. Hybrids were planted (May 4) into a randomized complete block design, with each hybrid by irrigation treatment being replicated four times. Plot size was 42 m by 9 m wide (12 rows; 0.76 m spacing). Plant spacing did not differ between hybrids and treatments and was 85,500 plants ha^−1^. Weed and fertility management followed standard practices for the region and plants appeared to be free of both weeds and nutrient stress throughout the experiment. All plots received full irrigation until plants reached V7 (seven fully-expanded and “collared” leaves present) on June 26, after which irrigation was reduced in the dry treatment. The dry treatment was lifted again as the plants approached VT (anthesis) on July 27, but was implemented again once the plants achieved R3 (starch accumulating “milk” stage) on August 29. This was done to avoid stress through the most sensitive reproductive stages.

### Trait measurements

#### Biomass, grain yield, and leaf area

Shoot biomass samples were collected for all hybrids in both treatments (wet, dry) immediately after plants reached physiological maturity (September 25 for dry and October 10 for wet treatments). Five representative plants were harvested from each plot for biomass and grain yield, giving 20 total plants per hybrid by treatment combination. Leaves, stems, ears, and grain were dried to constant mass and weighted to the nearest 0.01 g. The fresh leaf area of each harvested plant was measured (prior to drying) using a leaf area meter (LI-3100C, LI-COR, Lincoln, Nebraska, USA). Yield stability was calculated as the ratio of grain yield in the dry vs the wet treatments.

#### Stomatal conductance

Stomatal conductance was measured on all hybrids in both treatments (wet, dry) using hand-held steady-state porometers (Model SC-1, Meter Group Inc., Pullman, Washington, USA) between July 10 and July 14, 2017. Stomatal conductance was measured by walking continuously through the field from 0900 to 1500 each day. Four plants were measured within a single plot before moving on to the next adjacent plot. Each plot was measured ca 5 times throughout each day, giving a total of ca 95 individual measurements for each hybrid by treatment combination. Diurnal stomatal conductance trajectories for each hybrid by treatment combination were fit with quadratic models using the ‘nlsLM’ function in the minpack.lm package developed for R (Elzhov et al. 2016). Fitted maximum and midday (1400) values of stomatal conductance were then extracted from the quadratic models.

#### Leaf water potential

Leaf water potential was measured on all hybrids in both treatments (wet, dry) at midday (1200-1400) and at predawn (0500-0630) using a Scholander pressure chamber (Model 3005, Soil Moisture Equipment Corp, Santa Barbara, California, USA) between July 17 and September 1, 2017. This sampling resulted in ca 10 midday and 33 predawn measurements for each hybrid by treatment combination.

#### Light-saturated net CO_2_ assimilation and stomatal response to VPD

These traits were measured only on hybrids in the wet treatment using two portable gas-exchange systems (Model LI-6400-40, LI-COR Biosciences, Lincoln, Nebraska, USA). Measurements were taken between June 27 and July 7, 2017. Briefly, each morning, one plant of each hybrid would be randomly selected from the field, severed at its base, wrapped in white plastic, and brought back to the laboratory, thus giving at least six replicates of each hybrid (~one of each hybrid per day). Plants were then re-cut under water (leaving the severed end submerged) in a climate-controlled room (temp = 25 °C; relative humidity > 60%). Light-saturated net CO_2_ assimilation rate was measured on the top-most, fully-expanded leaf under light saturated conditions (1800 μmol m-2 s-1). Chamber temperature and VPD were kept below 30 °C and 1.5 kPa for at least 20 minutes prior to recording maximal measurements. After maximal measurements were recorded, stomatal conductance was measured under increasing VPD, from 1.5 kPa to 3.0 kPa in 0.2 kPa steps. Stomatal response to VPD was mostly flat, with a slight decline between 2.5 kPa and 3.0 kPa. To quantify the change in slope between 2.5 kPa and 3.0 kPa, spline models were fit to g_S_~VPD data using the ‘loess’ function in R and differences in slope extracted from these fitted models.

#### Pressure-volume curves

Pressure-volume data were measured only on hybrids in the wet treatment. The theory and assumptions of pressure-volume data have been discussed at length elsewhere (Schulte and Hinckley 1985; Ding et al. 2014). Here, we report how we obtained the necessary data to build pressure-volume curves and the metrics we extracted from them. Pressure-volume data were measured on six days between August 8 and August 17, 2017. Briefly, on each day, one leaf from each hybrid was collected randomly from the wet treatment during predawn hours (0500 – 0600), immediately placed in a sealable plastic bag and brought back to the laboratory (six replicates per hybrid). Leaves were repeatedly weighted to the nearest 0.0001 g and their leaf water potential measured with a Scholander pressure chamber (Model 3005, Soil Moisture Equipment Corp, Santa Barbara, California, USA). At least nine pressure-volume points were obtained in this way for each leaf. The reciprocal of pressure (1/MPa) was plotted against one minus relative water content (1 – RWC). Cell wall elasticity (ԑ), water potential at turgor loss (π_tlp_), osmotic potential at full turgor (π_o_), and leaf capacitance (C_Leaf_) were extracted from each curve. Leaf capacitance was estimated from the initial slope of the pressure volume curve prior to turgor loss.

#### Maximal leaf hydraulic conductance and leaf hydraulic vulnerability

Maximal leaf hydraulic conductance (K_leaf_max_) and leaf hydraulic vulnerability were measured only on hybrids in the wet treatment using the Rehydration Kinetics Method (Brodribb and Holbrook 2003) between July 14 and July 28, 2017. Briefly, on each day, ca four plants of each hybrid were cut at the base in the field during predawn hours (0500 – 0600), immediately placed in a white plastic bag, and brought back to the laboratory, where they were re-cut under water (in a large bucket), leaving their canopies still wrapped in plastic. Plants were removed from the bucket and dried down to a range of water potentials between −0.3 MPa and −4.0 MPa under a box fan. Once a plant had dried down to the desired water potential, two adjacent leaves near the top of the canopy were cut off. The water potential of one leaf was immediately measured in a Scholander pressure chamber (Model 3005, Soil Moisture Equipment Corp, Santa Barbara, California, USA). The other leaf was recut underwater, and the cut end allowed to re-hydrate (underwater) between 5 and 20 seconds whilst illuminated (1500 μmol m^−2^ s^−1^ PPFD). The water potential of the “re-hydrated” leaf was then immediately measured in the pressure chamber. Leaf conductance (K_leaf_) was calculated from its change in water potential and the increase in water volume during re-hydration, after Brodribb and Holbrook (2003):

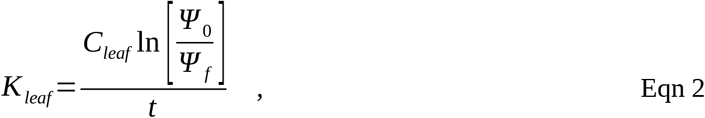

where K_leaf_ = the leaf conductance, C_Leaf_ = leaf capacitance, i.e., the change in leaf water content per change in water potential prior to turgor loss (obtained from the initial slope of pressure-volume curves), Ψ_0_ and Ψ_f_ are the leaf water potentials before re-hydration and after re-hydration, and t = the re-hydration time. K_leaf_ was measured in this way for at least 28 plants of each hybrid such that the decline in K_leaf_ could be plotted against leaf water potential to develop a “vulnerability curve”. K_leaf_~Ψ_leaf_ data were then fit with sigmoidal models after Pammenter and Vander Willigen (1998) using the ‘nlsLM’ function in the minpack.lm package developed for R (Elzhov et al. 2016). From this curve, we ranked the susceptibility of the hybrids to hydraulic failure according to their loss of K_leaf_ per unit decline in leaf water potential. For this purpose, we use the leaf water potential at which 50% of the maximal leaf conductance was lost (P_50_).

### Statistical analysis

All analyses, model fitting, and graphics were done in R 3.5.1. (R Core Team 2015). Bivariate correlation and multivariate analyses were done on hybrid mean data using the ‘lm’ function in base R and the ‘principal’ function in the ‘psych’ package for R, respectively. Given the small sample size (eight hybrids), bootstrapping was used to estimate the stability of fitted principal components and trait loadings (Babamoradi et al. 2013). If the standard deviation across bootstrapped samples was greater than 0.5 (range = −1 to +1), the loading was noted as “unstable”. Varimax rotation (an orthogonal method) was used to obtain more interpretable principal components and the ‘ggraph’ package for R was used to plot the results. The data used in this study are available in csv format (Appendix S1). All analyses and figures can be reproduced using these data. Additionally, the R code written to perform all analyses and figures are available from the first author upon request.

## Results

We present first the main axes of variation across the traits, i.e. trait groupings (Fig. 1). We note that because our emphasis was focused on performance under irrigated conditions, more traits were measured on fully irrigated plants (wet treatment) than on plants growing under water stress (dry treatment). Thus, these two data-sets (wet, dry) were analyzed independently.

**Figure 1.**
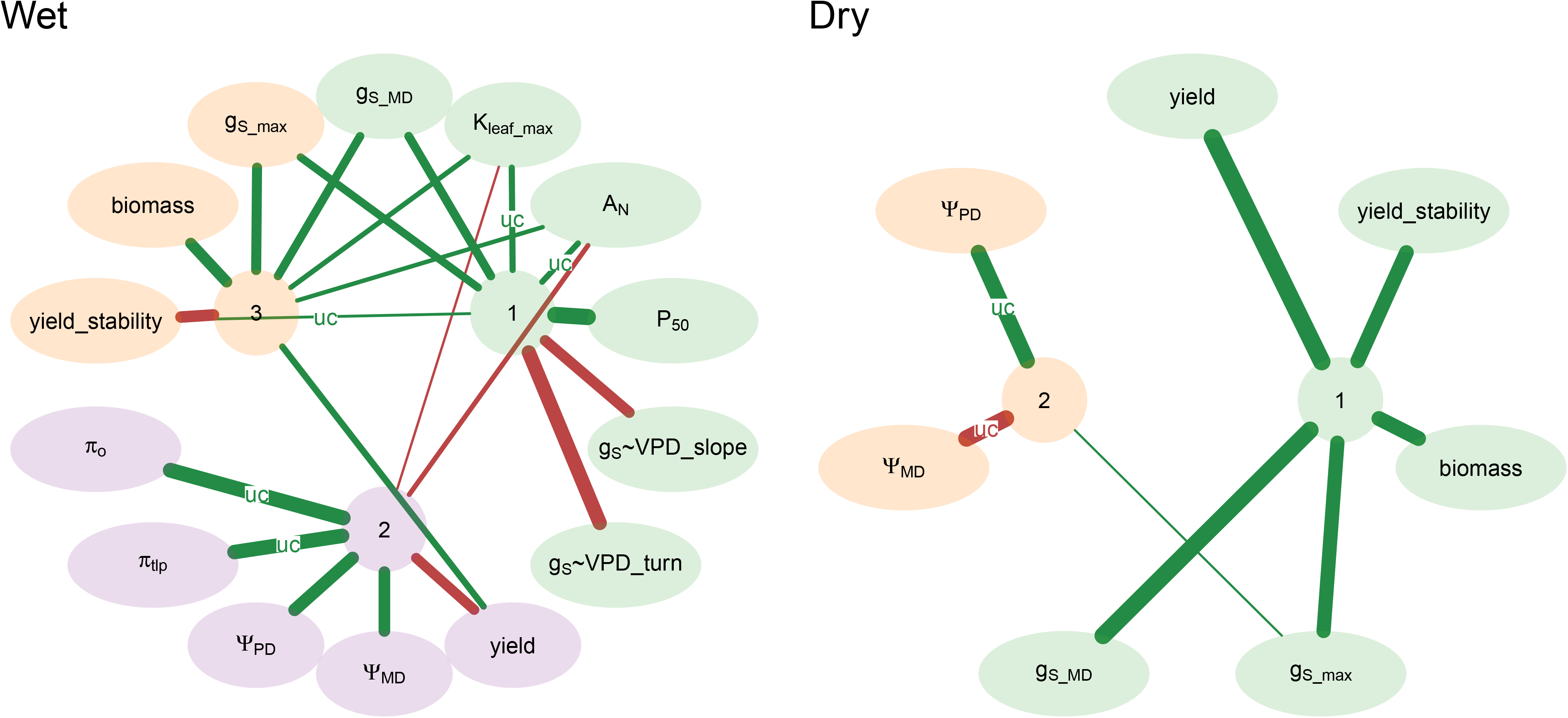
Orthogonality rotated principal components (colored circles) and trait groupings (colored ellipses). Positive loadings and negative loadings are denoted with green and red connections, with wider connections indicating larger standardized coefficients (0.5 to 1.0). Coefficients with standard deviations greater than 0.5 (range = −1 to +1) after bootstrapping are denoted as uncertain (UC) and should be interpreted with caution. Wet (fully-watered) and dry treatments (40% of reference crop ET) are shown in the left and right panels, respectively. g_S_max_ and g_S_MD_ = maximal and midday stomatal conductance (gas-phase water conductance), respectively. K_leaf_max_ = the maximal rate of leaf hydraulic conductance (liquid-phase water conductance). A_N_ = the light-saturated rate of net CO_2_ assimilation. P_50_ = the leaf water potential at which 50% of K_leaf_max_ was lost. g_S_ ~VPD_slope = the rate of stomatal conductance decline (mmol kPa^−1^) when the VPD was equal to 3.0 kPa, i.e. the first derivative of the g_S_~VPD function where VPD = 3. g_S_~VPD_turn = the VPD initiating stomatal closure. Yield = end-of-season grain yield. Ψ_MD_ and Ψ_PD_ = the leaf water potential during the middle of the day (1200-1400) and during predawn hours (0500-1630), respectively. π_tlp_ and π_o_ = the osmotic potential at turgor loss and at full turgor, respectively. Yield_stability is the ratio of grain yield produced in the dry treatment relative to that produced in the wet treatment. Biomass = the end-of-season biomass of all above-ground plant components (stems, leaves, reproductive structures).

Across all hybrids within the wet treatment, water transport capacity (K_leaf_max_), midday stomatal conductance (g_S_), and photosynthesis (A_N_) appeared bundled together as a single axes of variation (principal component 1; PC1) (Fig. 1 “Wet”). Beyond this, there was also strong alignment among maximal stomatal conductance (g_S_max_), growth (biomass increment), and grain yield, which manifested as a separate principal component (PC3), but with linkages to photosynthesis and water transport capacity (PC1) via midday and maximal stomatal conductance (Fig. 1 “Wet”). Interestingly, more negative hydraulic status, including lower osmotic adjustment (π_o_), leaf water potential at turgor loss (π_tlp_), and the operating water potentials during both midday (Ψ_MD_) and predawn hours (Ψ_MD_) were associated with greater yield along the second principle component (PC2), suggesting that higher yielding hybrids removed more water from the soil than poorer yielding hybrids (Fig. 1 “Wet”). Additionally, there was alignment between the VPD required to initiate stomatal closure (g_S_~VPD_turn), the rate of stomatal closure when VPD was equal to 3.0 kPa (g_S_~VPD_slope), and the first axis of variation (PC1; K_leaf_max_, g_S_MD_, and A_N_). The direction (inverse) and alignment of this variation suggests that early-closing (at low VPD) and fast-closing stomata (steeper g_S_~VPD slope) were associated with higher midday stomatal conductance, higher leaf conductance, and greater CO_2_ assimilation (Fig. 1 “Wet”). Also in alignment with this axis (PC1) was the rate at which leaf conductance was lost (per unit Ψ_leaf_) (P_50_), such that hybrids with higher stomatal conductance at midday also had leaves that were more susceptible to hydraulic failure at a given water potential. Bootstrapping revealed that the small sample size (eight hybrids per trait) resulted in relatively unstable loadings for yield stability (with PC1), K_leaf_max_ and A_N_ (PC1), π_o_ and π_o_ (PC2) in the wet treatment, and water potential (Ψ_PD_, Ψ_MD_) (PC2) in the dry treatment. This means that some hybrids exhibited specifically high leverage on the loading factor, and when they were removed from the analysis the value of the loading decreased meaningfully. As such, these linkages should be interpreted with caution.

An important difference between the wet and dry treatments was the much stronger alignment among growth, grain yield, and midday stomatal conductance in the dry treatment (Fig. 1 “Dry”). Indeed, the bivariate correlations between midday stomatal conductance and biomass (r^2^ = 0.69; p = 0.010) and grain yield (r^2^ = 0.91; p < 0.001) were markedly high in the dry treatment (Fig. 2 a,b), suggesting very close coordination between the achievable stomatal conductance in the middle of the day and growth in the dry treatment (Fig. 2B). Midday stomatal conductance was also strongly correlated with light-saturated net CO_2_ assimilation (symbol size) in the wet treatment (r^2^ = 0.58; p = 0.029) (Fig. 2B, 3A).

**Figure 2.**
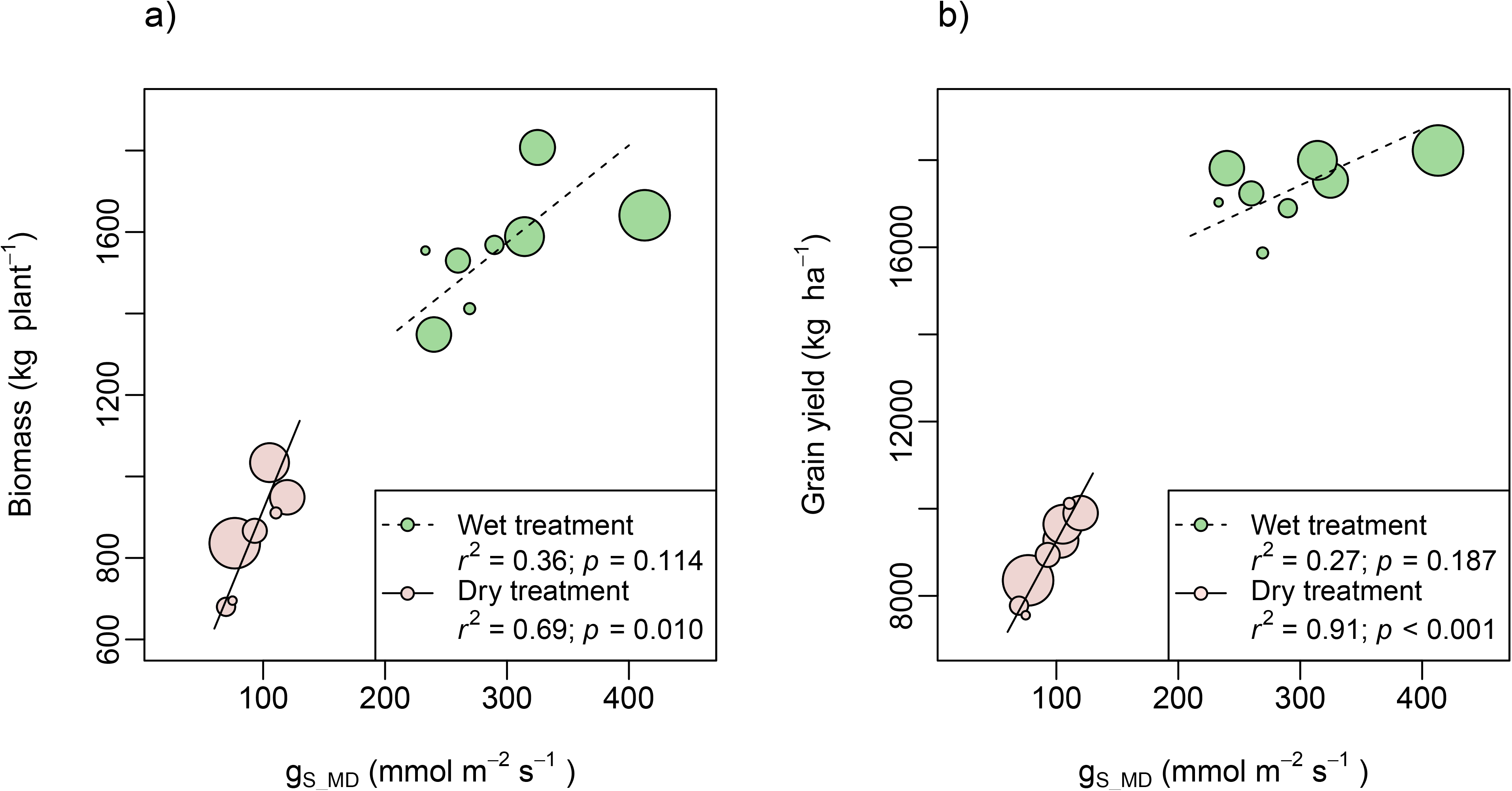
Bivariate plots of the linkages between grain yield and midday stomatal conductance (g_S_MD_) (a), and between biomass and midday stomatal conductance (b). Each symbol represents a single hybrid mean value (biomass and grain yield; n=4). Symbol size has been scaled to the light-saturated rate of net CO_2_ assimilation.

Although the bivariate models generally supported the multivariate analyses, there were a few differences worth noting. Specifically, there were strong correlations among traits that in some cases loaded on different principal components. For example, although the leaf’s capacity for water transport (K_leaf_max_) and the stomatal conductance at miday (g_S_MD_) loaded primarily on PC1, these traits also correlated strongly with plant growth (biomass increment) and grain yield (components of PC2 and PC3) (Fig. 3 “Wet”). This suggests strong coordination between the liquid water conductance, gas-phase conductance, net CO_2_ assimilation, and growth, but especially in the dry treatment (Fig. 3 “Dry”). The PCA and bivariate results suggest close linkage between the osmotic potential (π_o_) and grain yield, that is not likely manifesting through either A_N_ or biomass increment (weak linkages), suggesting a more proximal relationship between these traits (Fig. 3 “Wet”).

**Figure 3.**
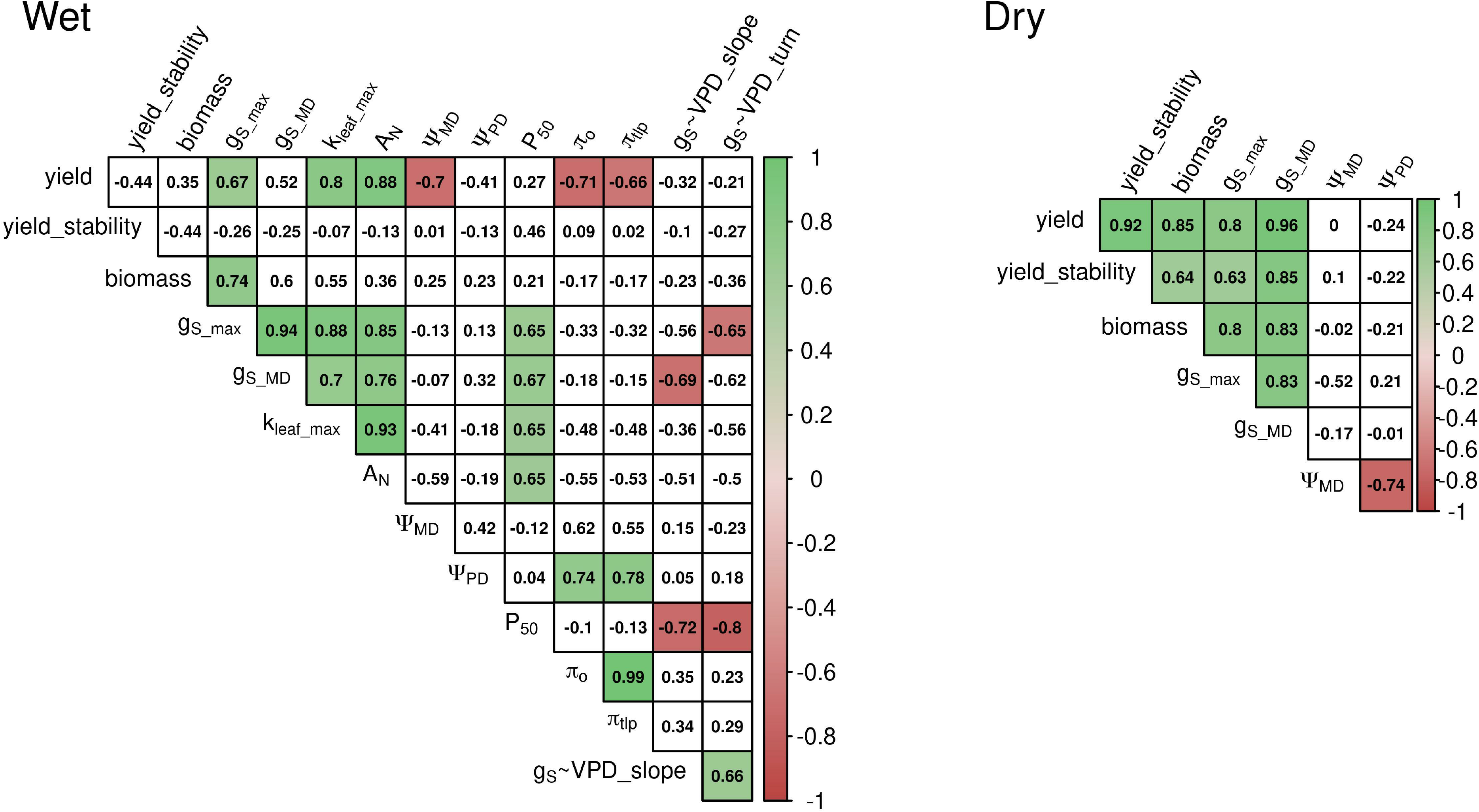
Correlation matrices for both wet (left panel) and dry (right panel) treatments. Correlation coefficients are denoted by text and by color, with increasing color intensity indicating increasing correlation strength. Variable descriptions are the same as given in Figure 1.

Considering both the multivariate and bivariate analyses, there appears to be strong and logical linkages that exist across hybrids. Although the strength of these linkages shift somewhat between the wet and dry treatments, they appear to be underpinned by the same physiological processes, that is, the acquisition of water, its transport to the sites of evaporation/photosynthesis, gas-exchange, growth, and finally, grain yield.

## Discussion

### Traits conferring improved performance under wet and dry conditions

Taken together, the results of the multivariate and bivariate analyses suggest a logical and tightly coordinated bundle of traits (processes) leading to improved performance under wet and dry treatments. Importantly, even though fewer traits were measured in the dry treatment than in the wet treatment, the traits and linkages conferring better performance were similar in both cases, as has been reported before (Gleason et al. 2019). High positive correlation between maximal stomatal conductance and grain yield under both wet and dry conditions suggests that, at least under the soil and climate of the study site and across the hybrids examined here, the maintenance of stomatal conductance throughout the day appears to be required for supporting daily net biomass accumulation. This result is supported by previous efforts to understand whole-plant functioning in crops, as well as wild species, in that water transport (~K_leaf_) to the stomata (*g_S_*) drives gas-exchange (~A_N_), and therefore, improved growth and yield (Blum 2009; Brodribb et al. 2015; Gleason et al. 2017a, 2019; Xiong and Nadal 2020).

Given the apparent importance in maintaining water transport and stomatal conductance, we might also expect better performing hybrids to be more resistant to embolism (lower P_50_) than poorer performing hybrids (Ryu et al. 2016), but our results do not support this. Rather, hybrids that exhibited better performance (higher midday g_S_, K_leaf_, growth, yield) under both wet and dry conditions also exhibited larger reductions in hydraulic conductance at low water potential (i.e., they exhibited higher P_50_ values). This result is nearly identical to a previous experiment using the parents of the Nested Association Mapped population (Gleason et al. 2019), and suggests that the better performing (growth and yield) hybrids/inbreds in these studies achieved greater stomatal conductance and gas exchange, not by having vasculature and/or stomata that are less sensitive to low water potential, but rather, by having higher hydraulic and stomatal conductance in the first place. For example, even though the hybrids/inbreds that “win the race” had more sensitive stomata and more vulnerable hydraulic pathways, they were still able to achieve greater liquid and gas-phase conductances through the middle of the day.

There are two trait combinations that, in theory, will lead to a higher sustained K_leaf_ during late morning and midday hours. Firstly, xylem that is more embolism resistant (lower P_50_) will exhibit smaller reductions in K_leaf_ as water potential declines through the day. Secondly, for a given embolism resistance, plants can start out in the morning with higher K_leaf_, such that their K_leaf_ remains sufficiently high during the day. As such, these two traits (higher K_leaf_ vs lower P_50_) represent functionally equivalent strategies. It is interesting to note then that either natural selection, artificial selection, or both have appeared to favor hybrids (this study) and inbreds (Gleason et al. 2019) with higher K_leaf_, rather than lower P_50_. Given that this result has now been reported in two independent maize experiments using different populations, it may be worth considering the relative costs and risks of these two alternative strategies. It is also noteworthy that high K_leaf_ would likely only be beneficial if embolism in maize is reversible at night via root pressure, which has been reported previously in this species (Steudle et al. 1987; Gleason et al. 2017b).

Osmotic adjustment in the leaves of the hybrids examined here was closely correlated with end-of-season grain production, pre-dawn water potential, and marginally with water transport and photosynthesis, again, suggesting a logical network of traits leading to improved performance (Figs. 1 & 3, “wet”). Although osmotic adjustment is a known beneficial drought response in vascular species, including maize (Ashwini et al. 2019; Beseli et al. 2019), our results here suggest that it might serve as a more effective breeding target if other closely aligned traits, namely K_leaf_ and A_N_ can also be targeted.

### Water transport vs transpiration efficiency

It is clear from our current understanding of plant physiology and drought that the improvement of individual plant traits in isolation of one another will not result in the best crop performance outcomes. Although the results here suggest that higher water transport efficiency might be a good strategy for water limited environments, there is evidence that traits conferring higher transpiration efficiency might also confer enhanced performance under similar conditions (Zaman-Allah et al. 2011; Vadez et al. 2014; Messina et al. 2015; Sinclair et al. 2017). It is possibly that different strategies may produce advantages depending on the availability of soil water across time and space, as well as the balance between atmospheric and soil aridity (Eqn. 1). Although this idea has yet to be tested rigorously, it should be understood that any attempt to increase either instantaneous or seasonally-integrated transpiration efficiency by reducing transpiration (even in the middle of the day) would have likely reduced the performance of the hybrids we evaluated here, as well as elsewhere (Jordan et al. 1983; Gowda et al. 2011; Gleason et al. 2019; Palta and Turner 2019). In our view, the efficacy of both of these broad crop strategies (water extraction and transport, transpiration efficiency) to confer improved performance under drought are supported by sound theoretical principles, and as such, this important research question remains low-hanging fruit for both experimentalists and modelers.

## Conclusions

Given that photosynthesis relies directly on water transport and stomatal conductance, we suggest that these traits could, in theory, be manipulated to improve crop species. However, this remains a difficult task for several reasons. Firstly, xylem traits are time-consuming, expensive, and require specific expertise to measure, and are therefore currently not well suited for high through-put methods. Additionally, the field of hydraulic physiology is relatively new, and recent advances in this field have not yet been transferred to other disciplines. However, even given these difficulties and the narrow scope of the present study (eight hybrids grown at a single site), the results we report here suggest that xylem is a common failure point in the water transport pathway (soil water --> xylem --> gas-exchange), and its performance correlates strongly with both gas exchange and growth (Brodribb 2009; Gleason et al. 2017a; Martin-StPaul et al. 2017; Xiong and Nadal 2020). As such, we suggest that xylem functioning, as well as the regulation and loss of conductance both within and outside the xylem (Scoffoni et al. 2017; Xiong and Nadal 2020), might be good candidates for breeding programs if these traits can be measured quickly and at the appropriate scale. Furthermore, considering traits as connected networks that manifest as effective crop strategies, as done here across a small group of hybrids, can help to identify novel avenues for crop improvement.

## Abbreviations

Ψ_MD_: leaf water potential during midday hours (1200 - 1400 hrs)
Ψ_PD_: leaf water potential during predawn hours (0500 - 0630 hrs)
Ψ_S_, Ψ_L_: water potential of soil and leaf, respectively
π_o_: leaf osmotic potential at full turgor
π_tlp_: leaf osmotic potential at turgor loss
ε: cell wall modulus of elasticity
L_A_: leaf area
A_N_: light-saturated, net CO_2_ assimilation rate
D: the leaf-to-atmosphere vapor pressure deficit
E: transpiration
g_S_: stomatal conductance
g_S_max_: maximum achievable stomatal conductance
g_S_MD_: stomatal conductance during the middle of the day (1400 hrs)
g_S_~VPD_slope: slope of the g_S_~VPD function when VPD is equal to 3.0 kPa
g_S_~VPD_turn: VPD value where the slope of the g_S_~VPD function becomes negative
k_leaf_max_: maximal leaf-specific hydraulic conductance
K_x_: xylem-specific conductivity
L: path-length between soil water and the sites of evaporation within the leaf
P_50_: leaf water potential resulting in a 50% loss of maximal leaf conductance

## Author Contributions

SMG and LHC developed the original idea and design for the experiment. SMG did the analyses and wrote the first manuscript draft. SMG, LHC, LN, and CH performed the trait measurements. RB and SC contributed germplasm and assisted with experimental design. All authors contributed equally to manuscript revisions.

## Acknowledgments

We warmly thank the scientists, technicians, collaborators and students that assisted with this study. We would like to especially thank Nora Flynn, Ryan Barton, Kelly Nelson, Jerry Buchleiter, Ross Stewart, and Garrett Banks for their kind assistance and enthusiastic support in the field.

## Supporting information

**S1** Mean values and standard deviations for all hybrid and treatment combinations (gleason_et_al.csv)

## Data availability statement

The data that supports the findings of this study are available in the supplementary material of this article

